# Pin1-promoted SUMOylation of RNF168 restrains its chromatin accumulation

**DOI:** 10.1101/2022.03.22.485326

**Authors:** Anoop Singh Chauhan, Alexander J. Garvin, Mohammed Jamshad, Joanna R. Morris

## Abstract

The E3 ubiquitin ligase RNF168 is a rate-limiting component of DNA double-strand break signalling that acts to amplify histone ubiquitylation. The confining of RNF168 chromatin spreading around DNA break sites has been related to constraining its expression levels and to the removal of ubiquitin-conjugates to which it binds. Here we identify a new mechanism that suppresses RNF168 amplification at chromatin. We show that depletion of Peptidyl-prolyl cis-trans isomerase NIMA-interacting 1 (Pin1) or mutation of the Pin1-binding site of RNF168 increases DNA-damage site accumulation of RNF168 to supraphysiological levels without impacting RNF168 expression levels. Pin1 promotes SUMOylation of RNF168 by SUMO2/3, and a SUMO conjugation site on RNF168 is required to restrict RNF168 accumulations. We find Pin1-SUMO mediated-regulation of RNF168 is associated with cellular radioresistance. These data demonstrate that the regulation of chromatin ubiquitylation is actively suppressed by modification of RNF168 to counteract excessive RNF168-ubiquitin spreading.

## Introduction

The signaling cascade activated by DNA double-strand breaks (DSBs) depends on several post-translational modifications orchestrated by phosphorylation and ubiquitylation. Ubiquitination is driven by a series of E3 ubiquitin (Ub) ligases, begun by the chromatin modification activities of RNF8 and RNF168 (Doil et al. 2009; Huen et al. 2007; Kolas et al. 2007; Mailand et al. 2007; Stewart et al. 2009). RNF8 catalyses K63 linked poly-ubiquitylation of Histone H1 (Thorslund et al. 2015) and L3MBTL2 (Nowsheen et al. 2018) and the ubiquitin-binding domains of RNF168 then mediate its recruitment to these sites (Doil et al. 2009; Stewart et al. 2009; Takahashi et al. 2018). RNF168 in turn, catalyses ubiquitylation of Histone H2A and H2A variants (Kelliher et al. 2020; Mattiroli et al. 2012). RNF168 mediated ubiquitylation generates a positive feedback loop through its recognition of autocatalysed ubiquitylated H2A (Kelliher, Ghosal, and Leung 2021). Modified H2A is also bound by 53BP1 (Fradet-Turcotte et al. 2013) and BARD1-BRCA1 (Becker et al. 2021), which in turn support non-homologous end-joining DNA repair or homologous recombination, respectively (reviewed in (Schwertman, Bekker-Jensen, and Mailand 2016)).

While RNF168 is a fulcrum of the ubiquitin-mediated damage response, it is capable of run-away positive feedback. Loss of proteins that restrict RNF168 expression levels (TRIP12/UBR5) or increased RNF168 expression through chromosome amplification results in an exaggerated signalling response deregulated sequestration of downstream proteins, altered DSB repair and increased local transcriptional silencing (Chroma et al. 2017; Gudjonsson et al. 2012). RNF168 is a short-lived protein and is stabilised following DNA damage by the action of deubiquitylating enzymes (DUBs) such as USP7 and USP34 (Sy et al. 2013; Zhu et al. 2015). In addition, to RNF168 mediated catalysis, chromatin ubiquitylation levels are maintained by the DUB activity of USP3 and USP16 (Nicassio et al. 2007; Zhang, Yang, and Wang 2014).

The phosphorylation-dependent peptidyl-prolyl isomerase (PPAIse), Pin1, regulates CtIP and BRCA1 turnover (Liou, Zhou, and Lu 2011; Steger et al. 2013), promotes BRCA1-BARD1 interaction with RAD51 during replication stress (Daza-Martin et al. 2019) and its WW domain can purify several proteins critical to DNA double-strand break repair (Steger et al. 2013). Here, we show that Pin1 suppresses excessive chromatin ubiquitylation at sites of DSBs through direct regulation of RNF168, suppressing its unrestricted spreading. Pin1 interaction regulates RNF168 SUMOylation, which contributes to the suppression of RNF168 accumulations. Unrestricted accumulation of RNF168 is reflected in increased 53BP1 accumulation and correlates with increased sensitivity of cells to ionising radiation. Together these results uncover a surprising mechanism that cells use to restrict the extent of chromatin ubiquitylation in response to DNA damage.

## Results

### Potential regulation of RNF168 on chromatin by Pin1

As RNF168 can bind to and amplify ubiquitin conjugates generated by its own activity, we hypothesised that mechanisms beyond the suppression of its expression and chromatin de-ubiquitination might be needed to contain its spread. We considered that direct post-translational modification (PTMs) might be associated with this activity and mined publicly available “eukaryotic linear motif (ELM) resource” (Kumar et al. 2020) for potential RNF168 motifs. Intriguingly, we found the presence of numerous potential Pin1 binding sites both within and near the ubiquitin (Ub)-dependent DSB recruitment modules (UDMs) of RNF168 (Supplementary figure 1A).

We examined the accumulation of endogenous RNF168 following Pin1 depletion in cells treated with ionising radiation (IR) and found increased intensity of RNF168 at sites decorated with γ-H2AX (Figure 1A and B). This observation was also recapitulated in cells expressing exogenous RNF168 in both untreated and IR exposed conditions (Figure 1D-G). It was noted that in untreated cells, the spread of accumulated RNF168 was beyond the boundaries of DNA lesions marked by γ-H2AX (Figure 1D, right panel, Supplementary figure 1B). Consistent with these observations, we found increased RNF168 protein levels enriched on chromatin in Pin1 depleted cells (Figure 1H). These changes appeared unrelated to total RNF168 protein levels which were unchanged in Pin1 depleted cells (Figure 1H).

**Figure 1.**
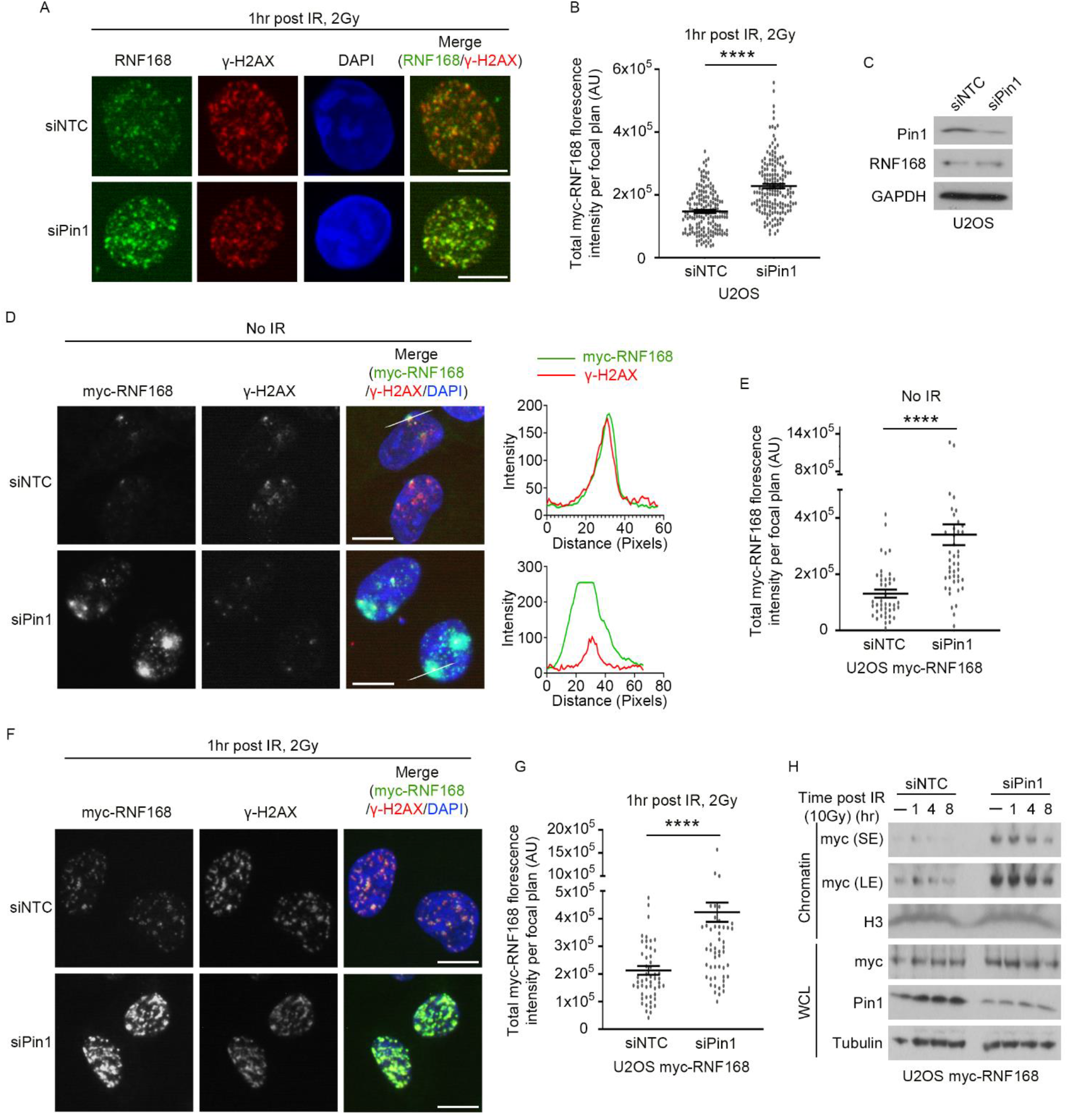
Pin1 suppresses RNF168 chromatin localisation. A. Representative images of RNF168 foci after treatment with ionising radiation (IR). U2OS cells were transfected with non-targeting control (NTC) or Pin1 siRNA for 72 hrs and treated with IR, 2 Gy. 1 hr after IR cells were fixed and stained for endogenous RNF168 and γ-H2AX. Scale bars 10 μm. B. Quantification of RNF168 foci intensity from A. Data is mean ± s.e.m, n= 170 cells. p-value <0.0001. C. Western blot of Pin1 protein following treatment with non-targetting siRNA (siNTC) and Pin1 siRNA (siPin1). D. Representative images of exogenous RNF168 foci. U2OS cells stably expressing myc-RNF168 were treated with siNTC or siPin1 for 72 hrs and stained for myc and γ-H2AX. Scale bars 10 μm. The right panel shows the fluorescence intensity profiles along the line for myc and γ-H2AX. E. Quantification of myc-RNF168 foci intensity from D. Data is mean ± s.e.m, n= 42 cells for siNTC and 48 cells for siPin1. p-value <0.0001. F. Representative images of myc-RNF168 foci formation after IR. U2OS cells expressing myc-RNF168 were treated with indicated siRNAs and stained for myc and γ-H2AX after 1 hr post IR, 2 Gy. Scale bars 10 μm. G. Quantification of myc-RNF168 foci intensity from F. Data is mean ± s.e.m, n= 54 cells for siNTC and 66 cells for siPin1. p-value <0.0001. H. Western blot of chromatin fraction or whole cell lysate (WCL) for myc, histone H3, Pin1 and tubulin. U2OS cells expressing myc-RNF168 were treated with siNTC or siPin1 for 72 hrs. Cells were treated with 10 Gy of IR and collected after indicated time points to prepare chromatin fraction or WCL. SE: short exposure, LE: Long exposure.

### ResiduesThr208-Pro209 of RNF168 regulate its chromatin accumulation

Pin1 is unique among PPIases in that in addition to the catalytic domain that isomerises prolyl bonds between prolines and the preceding amino acid, it has an N-terminal “WW” domain (referring to two required Trp residues) that targets the enzyme to pSer/Thr-Pro motifs in substrates (Andreotti 2003; El Boustani et al. 2018; Liou, Zhou, and Lu 2011; Zhou and Lu 2016). We generated and purified the wild type (WT) GST-Pin1-WW domain and a mutant form, GST-WW-W34A, that is unable to bind phosphorylated Serine or Threonine (Lu et al. 1999). The WT-WW domain but not the mutant efficiently co-precipitated RNF168 from U2OS and HeLa cells (Figure 2A & B). Moreover, we found RNF168 was more enriched by the Pin1-WW domain from lysates of cells treated with ionizing radiation (IR) than from untreated cells (Figure 2C). These data point to potential direct regulation by Pin1.

**Figure 2.**
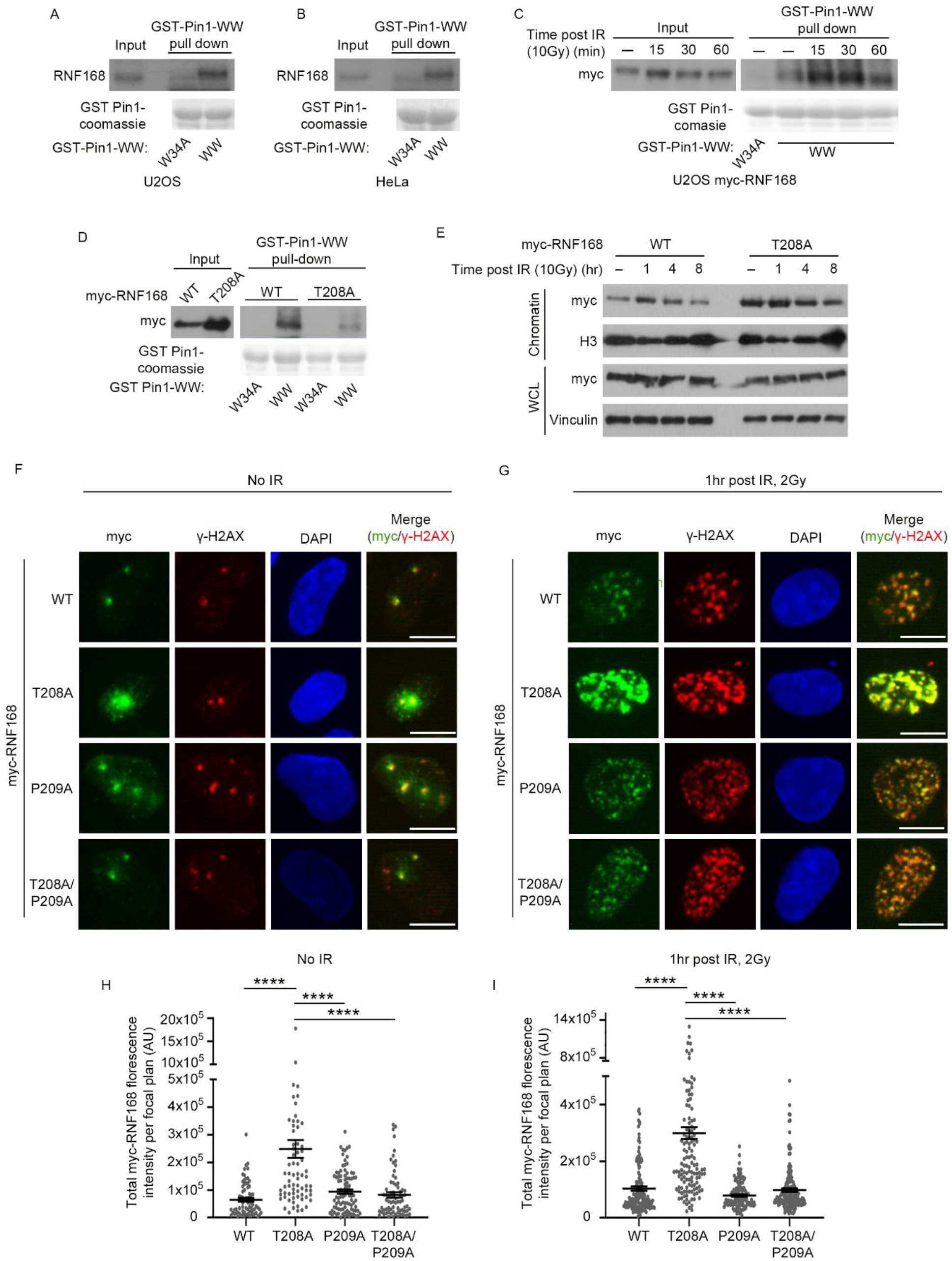
Pin1 regulate RNF168 accumulation by binding to Thr208/Pro209 motif. A. Pull down of endogenous RNF168 by GST-fused WW or W34A mutant domain of Pin1 from U2OS cells. B. Pull down of endogenous RNF168 by GST-fused WW or W34A mutant domain of Pin1 from HeLa cells. C. Pull down of myc-RNF168 by GST-fused WW or W34A mutant domain of Pin1 from U2OS cells treated with 10Gy of IR. Cell lysate were made at indicated time points after IR and subjected for WW domain pull-down. D. U2OS cells expressing myc-RNF168 wild type (WT) or T208A mutant were subjected to pull down by GST-fused-WW or W34A domain of Pin1. E. Western blot of chromatin fraction or WCL probed for myc, histone H3 and vinculin. U2OS cells were treated with siRNAs targeting RNF168 for 72 hrs to deplete endogenous RNF168. Cells were simultaneously treated with 1 μg/ml doxycycline to induce expression of siRNA resistant myc-RNF168-WT or T208A mutant. Chromatin fraction and WCL were prepared following treatment with 10 Gy of IR at indicated time points. F. Representative images of foci formation of exogenously expressed RNF168 mutants. U2OS cells were treated with 1 μg/ml doxycycline for 72 hrs to induce expression of siRNA resistant myc-RNF168-WT, T208A, P209A and T208A/P209A mutants. Endogenous RNF168 was depleted simultaneously by treatment with siRNF168. Cells were fixed and stained for myc and γ-H2AX. Scale bars 10 μm. G. As in F, cells expressing RNF168-WT, T208A, P209A, T208A/P209A mutants were treated with 2 Gy of IR and fixed after 1hr. Cells were stained for myc and γ-H2AX. Scale bars 10 μm. H. Quantification of myc-RNF168 mutant foci intensity from F. Data is mean ± s.e.m, n= 73 for WT, 75 for T208A, 92 for P209A and 73 for T208A/P209A. p-value <0.0001 (****). I. Quantification of myc-RNF168 mutant foci intensity from G. Data is mean ± s.e.m, n= 152 for WT, 142 for T208A, 130 for P209A and 163 for T208A/P209A. p-value <0.0001 (****).

We next tested whether any of the three potential Pin1 binding motifs (Ser197/Pro198, Thr208/Pro209 or Thr230/Pro231) near the RNF168 UDM1 are required for WW-domain enrichment. We mutated each Ser/Thr to Ala and found T208A, but not S197A or T230A mutated RNF168 showed compromised interaction with the GST-WW-Pin1 domain (Figure 2D, Supplementary figure 2A). Moreover, T208A-RNF168 formed brighter IR-induced foci (IRIF) and showed greater chromatin enrichment than WT-RNF168, S197A-RNF168 or T230A-RNF168 (Figure 2E, Supplementary figure 2B & C). T208A-RNF168 showed no more total protein expression than WT-RNF168 (Figure 2E). Together these data correlate an interaction of Pin1 with Thr208/Pro209 of RNF168 and with altered chromatin localisation of RNF168.

In polypeptide chains, the majority of the peptide bonds present between amino acids adopt a *trans* conformation (Joseph, Srinivasan, and de Brevern 2012). Proline has a unique side chain, forming a five-membered ring, allowing both *cis* and *trans* prolyl bonds (MacArthur and Thornton 1991). To explore whether isomerisation of RNF168 at the pThr208/Pro209 site might be significant to its chromatin localisation, we mutated proline-209 to alanine, reasoning that the substitution increases the probability of *trans* isomer (Nakamura et al. 2012; Zhou et al. 2000). We compared the localisation intensity and chromatin enrichment of P209A-RNF168 and the double mutant, P209A-T208A-RNF168, with that of WT-RNF168 and T208A-RNF168. P209A-RNF168 showed levels of each that were comparable to WT-RNF168, remarkably, so too did the double mutant P209A-T208A-RNF168 (Figure 2F-I), indicating the deleterious impact of T208A was negated by P209A. These findings suggest presenting *trans* favoured conformation by-passes the need for T208 in suppressing excessive chromatin accumulation. We also noted that the T208A-RNF168 mobilization to DSBs was dependent on RNF8 (Supplementary figure 2D & E).

### Thr208- RNF168 restrains chromatin ubiquitylation

To examine the impact of excessive RNF168 chromatin accumulation mediated by Thr208-Pro209, we assessed chromatin ubiquitination and found that cells complemented with the T208A-RNF168 showed increased mono and di ubiquitylation of γ-H2AX after IR compared to cells expressing wild type RNF168 (Figure 3A). We also observed an increase in 53BP1 intensity at sites of γ-H2AX decorated chromatin in undamaged and irradiated T208A-RNF168 complemented cells (Figure 3B & C), consistent with the spreading of RNF168 observed (Figure 1D & Figure 2 F). These data suggest the increased accumulation of RNF168 results in increased ubiquitin signaling and 53BP1 recruitment.

**Figure 3.**
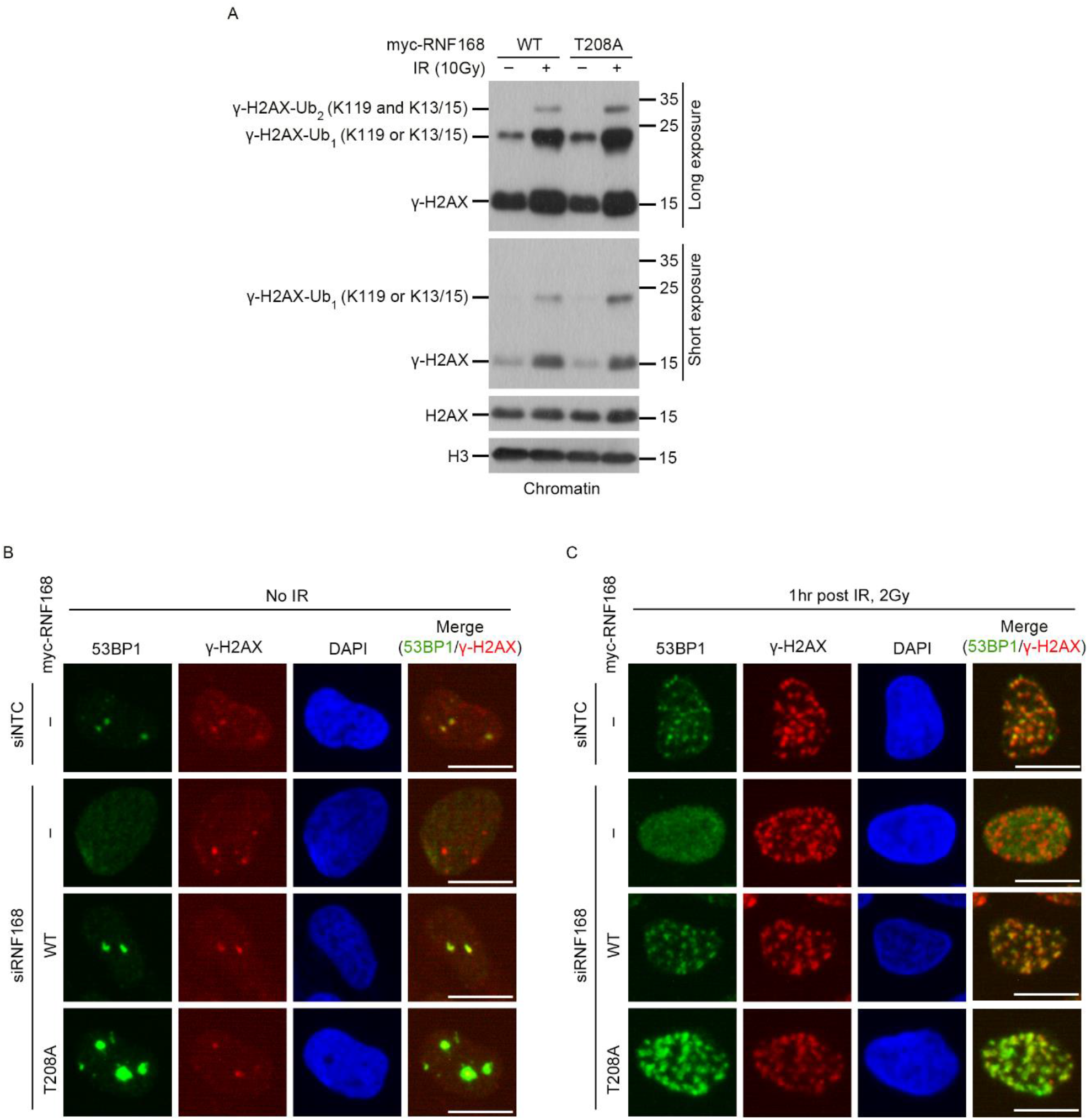
Pin1 binding deficient RNF168 promotes chromatin ubiquitylation. A. Western blot of chromatin fraction for histone γ-H2AX, H2AX and H3. U2OS cells were treated 1 μg/ml doxycycline to induce expression of myc-RNF168-WT or T208A mutant for 72 hrs. Endogenous RNF168 was depleted by treatment with siRNA. Later cells were treated with 10 Gy of IR and chromatin fraction was prepared for untreated or IR treated cells after 1 hr. B. U2OS were treated with siRNF168 for 72 hrs and incubated with 1μg/ml doxycycline to induce expression of siRNA resistant myc-RNF168-WT or T208A mutant. Cells were then fixed and stained for 53BP1 and γ-H2AX. Scale bars 10 μm. C. As in B, RNF168 knock down cells and cells expressing WT or T208A mutant myc-RNF168 were treated with 2 Gy of IR. After 1 hr cells were fixed and stained for 53BP1 and γ-H2AX. Scale bars 10 μm.

### Pin1 interaction with RNF168 is linked to its SUMOylation

We noticed that the Pin1 binding site at Thr208-Pro209 is adjacent to a previously described SUMO modification site at lysine 210 “SDPVtPkSEKKSKN” (Hendriks et al. 2014; Hendriks et al. 2017). To assess whether SUMOylation has the potential to impact RNF168 chromatin accumulation we depleted SUMO1 and SUMO2 with SUMO3 (SUMO2 and SUMO3 isoforms share 97% sequence identity). We found that depletion of SUMO2/3, but not SUMO1 increased RNF168 accumulation to γ-H2AX in undamaged and IR-treated cells and saw SUMO2/3 depletion increased RNF168 chromatin enrichment compared to control siRNA-treated cells (Figure 4A-F). We next examined if K210 of RNF168 is a SUMO2 modification site. We expressed WT-RNF168 and K210R-RNF168 with His-SUMO2 and nickel purified SUMO conjugates under denaturing conditions. WT-RNF168 was enriched and showed a band-size consistent with mono-SUMOylation; in contrast, K210R-RNF168 was not enriched (Figure 4G). We also noted that K210R-RNF168 showed greater chromatin enrichment than WT-RNF168, without exhibiting increased protein expression levels (Figure 4H). Taken together, these data suggest SUMOylation at RNF168 Lysine 210 also suppresses RNF168 chromatin spreading.

**Figure 4.**
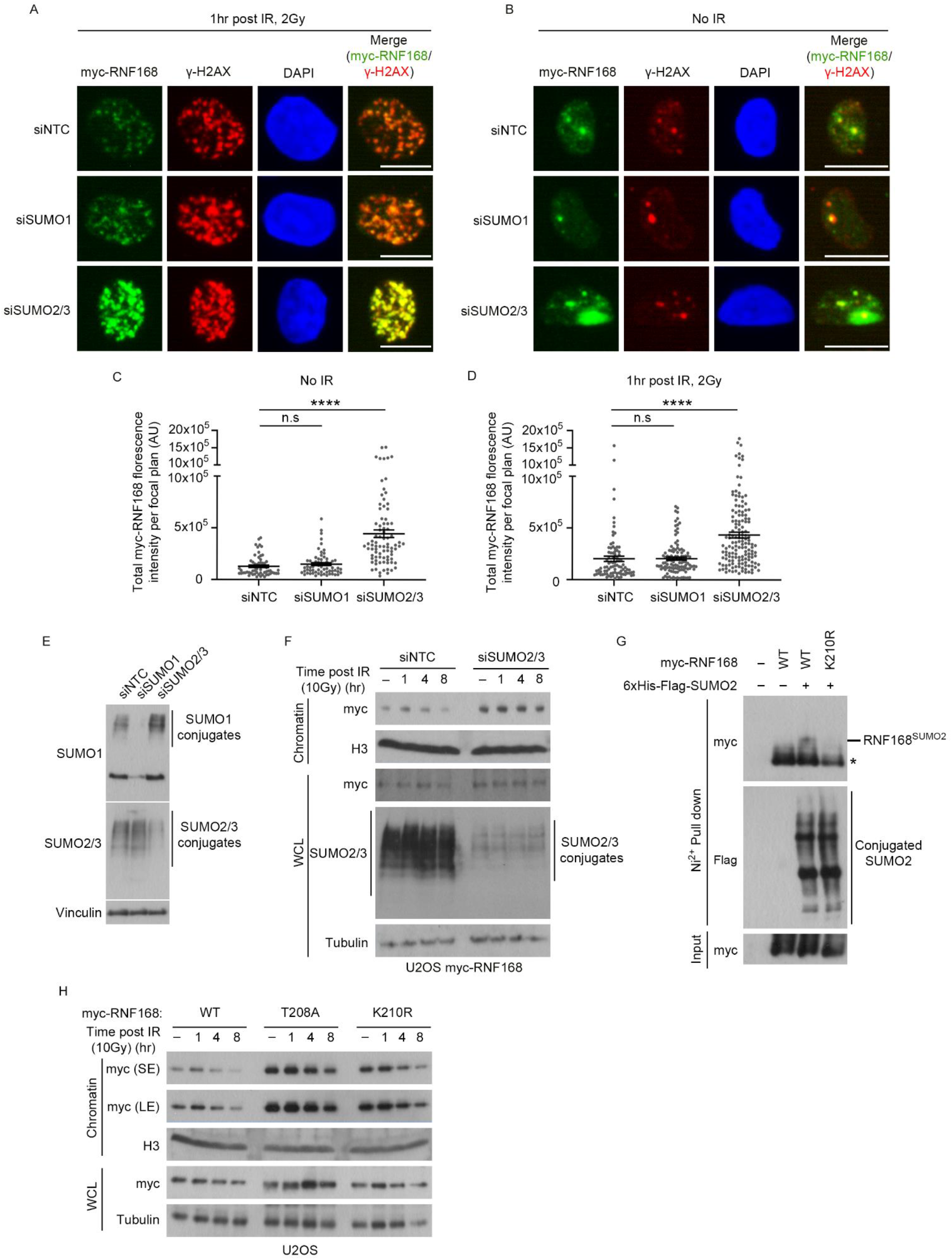
RNF168 SUMOylation by SUMO2/3 regulates its accumulation. A. U2OS cells were treated with indicated siRNAs for 72 hrs. Expression of myc-RNF168 was induced by the addition of 1 μg/ml doxycycline. Cells were then fixed and stained for myc and γ-H2AX. Scale bars 10 μm. B. As in A, after treatment with indicated siRNAs and doxycycline for 72 hrs, cells were treated with 2 Gy of IR. After 1 hr, cells were fixed and stained for myc and γ-H2AX. Scale bars 10 μm. C. Quantification of myc-RNF168 foci intensity from A. Data is mean ± s.e.m, n= 61 cells for siNTC, 65 cells for siSUMO1 and 84 cells for siSUMO2/3. p-value <0.0001 (****), n.s: not significant. D. Quantification of myc-RNF168 foci intensity from B. Data is mean ± s.e.m, n= 90 cells for siNTC, 110 cells for siSUMO1 and 146 cells for siSUMO2/3. p-value <0.0001 (****), n.s: not significant. E. Western blot to show depletion of SUMO1 and SUMO2/3 conjugates for D, E. F. Western blot of chromatin fraction and WCL for myc, histone H3, SUMO2/3 and tubulin. U2OS cells were treated with siNTC or siSUMO2/3 and 1 μg/ml doxycycline to induce expression of myc-RNF168 for 72 hrs. Later cells were treated with 10 Gy of IR and collected at indicated time points. G. HEK293 cells complemented with myc-RNF168-WT or K210R mutant were transfected with 6xHis-Flag-SUMO2. Cells were treated with 1μg/ml doxycycline for 72 hrs and additional 6 hrs with 10 μM MG132. SUMO2 conjugated proteins were enriched by His-Mag Sepharose Ni beads (Ni^2+^ pull down) under denaturing conditions and detected by western blotting. * = non-specific band. H. Western blot of chromatin fraction and WCL for myc, histone H3 and tubulin. U2OS cells were treated 1 μg/ml doxycycline to induce expression of WT, T208A and K210R mutant myc-RNF168 for 72 hrs. Cells were simultaneously incubated with siRNF168 to deplete endogenous RNF168 levels. Later, cells were treated with 10 Gy of IR and collected at indicated time points to prepare chromatin and WCL fractions.

To explore whether SUMOylation and isomerisation of RNF168 are connected, we precipitated His-SUMO2 from Pin1 depleted cells. Compared to control shRNA-treated cells, we noted a decrease in RNF168 enrichment in His-SUMO2-conjugates when Pin1 was depleted (Figure 5A). Moreover, when we expressed RNF168 variants and tested precipitated His-SUMO2 for RNF168, we observed less enrichment for T208A-RNF168 mutant compared to WT-RNF168 or to the P209A-T208A-RNF168 double mutant (Figure 5B). These data indicate Pin1, and specifically, the T208-P209 site influences RNF168 SUMOylation.

**Figure 5.**
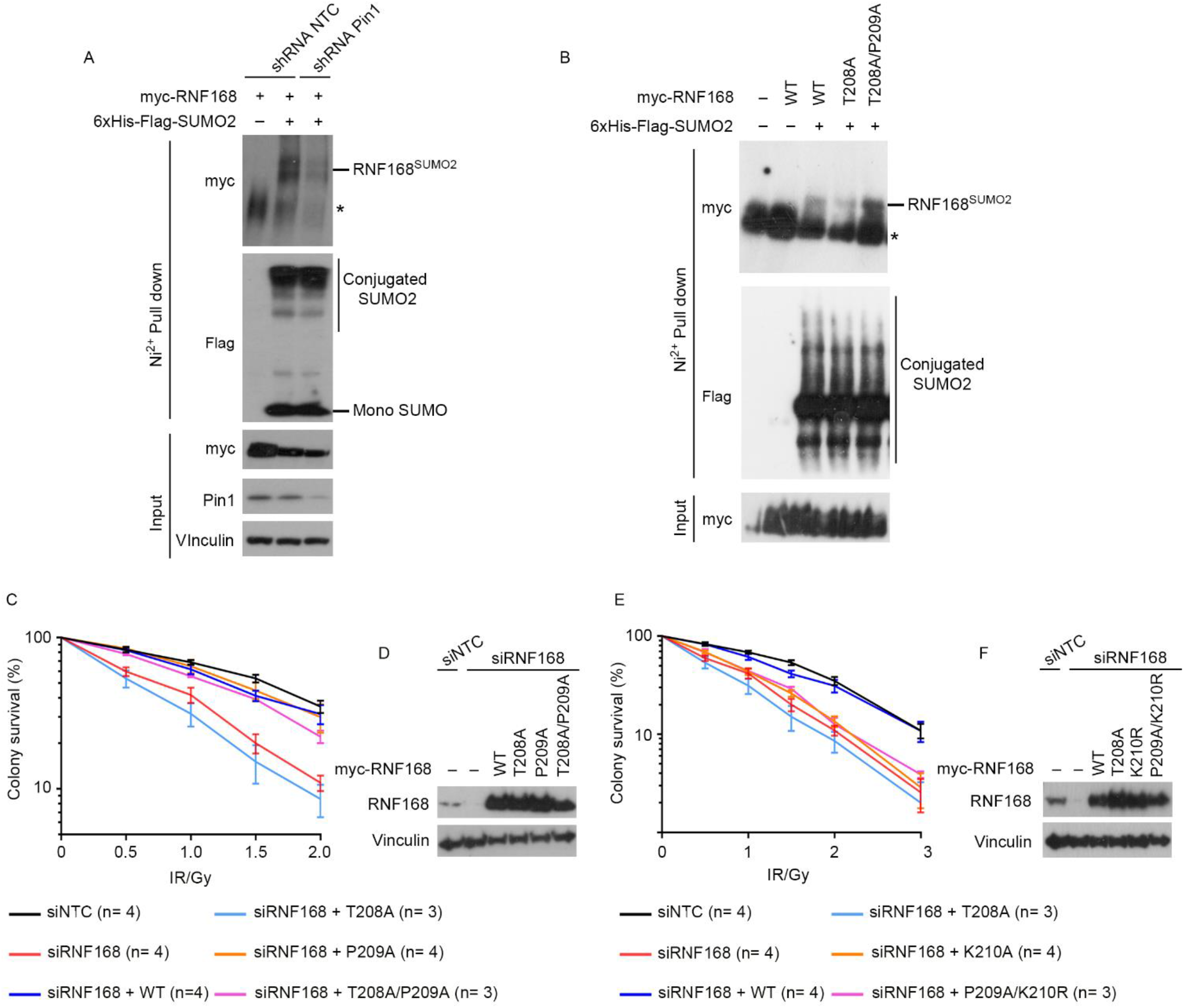
Pin1 regulates RNF168 SUMOylation and Pin1-SUMO sites support radioresistance. A. HEK293 cells containing shRNA control (NTC) or shPin1 were transfected with myc-RNF168 alone or in combination with 6xHis-Flag-SUMO2. Cells were treated with 1 mM IPTG for 72 hrs for Pin1 depletion. 10 μM MG132 was added for 6 hrs before lysate preparation. SUMO2 conjugated proteins were enriched by Ni^2+^ beads under denaturing conditions. * = non-specific band. B. HEK293 cells complemented with various RNF168 variants were transfected with 6xHis-Flag-SUMO2. Cells were treated with 1 μg/ml doxycycline for 72 hrs. Later SUMO conjugated proteins were pulled down with Ni^2+^ beads under denaturing conditions. SUMO conjugated RNF168 variants were detected by western blotting. * = non-specific band. C. Colony survival of U2OS cells depleted of RNF168 and complemented with WT, T208A, P209A and T208A/P209A variants of myc-RNF168 after treatment with indicated doses of IR. The number of replicates n for each condition is mentioned in parentheses. Data is mean ± s.e.m. D. Western blot to show depletion of RNF168 and complementation of RNF168 variants for C. E. Colony survival of U2OS cells depleted of RNF168 and complemented with WT, T208A, K210R and P209A/K210R variants of myc-RNF168 after treatment with indicated doses of IR. The number of replicates n for each condition is mentioned in parentheses. Data is mean ± s.e.m. F. Western blot to show depletion of RNF168 and complementation of RNF168 variants for E.

### RNF168-Pin1-SUMO site is critical for IR resistance

To test whether the RNF168 modification sites identified are relevant to cellular IR resistance, we examined colony survival of irradiated cells depleted for endogenous RNF168 and expressing siRNA-resistant variants. RNF168 depletion sensitised cells to IR, which could be rescued by complementation with WT-RNF168 (Figure 5C & D) but not T208A-RNF168. In contrast, cells complemented with T208A-P209A-RNF168 were not sensitive to IR (Figure 5C & D), consistent with the ability of the P209A to suppress the deleterious impact of T208A on RNF168 SUMOylation and on excessive chromatin accumulation.

We also tested the complementation of RNF168 depleted cells with K210R-RNF168 and with P209A-K201R-RNF168. Mutation of the lysine at 210 sensitised cells to IR (Figure 5E & F). However, the P209A-K210R-RNF168 mutant did not improve radioresistance over cells complemented with the K210R-RNF168 mutant (Figure 5E & F). These findings are in contrast to the results from cells expressing T208A/P209A-RNF168 over those expressing T208A-RNF168 and suggest the requirement for the K210 is dominant over the requirement for T208. Collectively, these data suggest the Pin1-SUMO site of RNF168 supports radiation tolerance.

## Discussion

Here we present the finding that the restriction of RNF168 chromatin accumulation and spreading requires a series of modifications of RNF168 itself. Suppression of Pin-WW domain interaction by the T208A-RNF168 mutation implies the T208 site is phosphorylated and pT208-RNF168 has been identified in bulk –phospho-proteome analysis (Koch et al. 2016). Our findings suggest RNF168 is modified at this site in response to IR, but the kinase responsible is currently unknown. Pin1-mediated change in protein conformation has been noted to increase the ubiquitylation of some substrates (Liou, Zhou, and Lu 2011; Orlicky et al. 2003) and recently to increase SUMO modifications (Zhang et al. 2020). Here we find a close association between Pin1-mediated conformational change of RNF168 and its SUMOylation. Indeed, comparative mutation analysis suggested that it is the SUMOylation, not the conformational change, that is most significant for confining RNF168 chromatin spreading. A study from the Mailand group found RNF168 modification by SUMO1 is mediated by PIAS4, which also promotes RNF168 expression (Danielsen et al. 2012), the SUMO ligase responsible for its SUMO2/3 modification at K210 may, or may not, be the same. SUMOylation of DNA-damage proteins has many roles but is most particularly linked to their clearance (Galanty et al. 2012; Garvin and Morris 2017; Garvin et al. 2019; Vyas et al. 2013; Yin et al. 2012). SUMO-mediated processing of RNF168 from chromatin would be one potential mechanism for how SUMOylation acts to restrict excessive RNF168 accumulations.

Our findings that RNF168 mutants with increased chromatin accumulation are associated with radio-sensitivity are in contrast with previous correlations between increased RNF168 spreading through increased protein levels, which showed increased NHEJ, expected to give radioresistance (Gudjonsson et al. 2012; Chroma et al. 2017). We speculate that RNF168 mutants may be unable to repair all of the breaks due to excessive spreading limiting the available RNF168, but do not discount other mechanisms, such as suppression of NHEJ by excessive 53BP1 recruitment.

Both Pin1 and the SUMO system are seen as pathways that might be targeted to improve the treatment of human cancers (Kroonen and Vertegaal 2021; Lu and Hunter 2014), and both are highly pleiotropic systems. Nevertheless, our findings are consistent with the requirement for SUMO and Pin1 in promoting radioresistance (Garvin and Morris 2017; Liu et al. 2019). In summary, we show that the positive feedback of Ub-binding and conjugation mediated by RNF168 at sites of damage is restrained by post-translational modifications of RNF168 itself.

## Materials and methods

### Cell culture and generation of inducible, stable cell lines

Parental Flip-In U2OS, HEK293 and HeLa cells were cultured in DMEM media (Sigma, RNBK7590) supplemented with 10% free fetal bovine serum (FBS) (Gibco, 10500-064) and 1% penicillin and streptomycin. Hoechst DNA staining was regularly performed to test for mycoplasma.

### Cloning, site-directed mutagenesis and primers

myc-tagged wild type RNF168 cloned into pCDNA5/FRT/TO plasmid was obtained from JRM laboratory (Garvin et al. 2019). Site-directed mutagenesis was performed in house by pfu DNA polymerase (Promega, M774A). Incorporated mutations were confirmed using sanger sequencing (Source Biosciences Nottingham). N-terminal 6xHis-Flag-tagged SUMO2 was cloned into BamH1 and Xho1 restriction sites of pcDNA5/FRT/TO plasmid by gene synthesis (GenScript Biotech). All primers used for site-directed mutagenesis are listed in **supplementary table 1**.

### Generation of inducible stable cell lines

Stable cells were generated by transfecting parental Flip-In U2OS and HEK293 cells with pCDNA5/FRT/TO plasmid containing myc-RNF168 wild type or its mutants along with Flp recombinase cDNA containing pOG44 plasmid in 3:1 ratio. Control cells were seeded alongside and transfected with pOG44 plasmid alone. After 48 hours, cells were selected by treatment with 100 μg/ml hygromycin (Thermo Fisher, 10687010). Expression of gene of interest was induced by treating cells with 1 μg/ml doxycycline (Merck, D9891) for 72 hours and confirmed by western blotting.

### Gene silencing and transfections

RNF168 and Pin1 knock down was performed by transfecting cells with siRNAs. siRNA transfections were carried out with Dharmafect1 (Dharmacon, T-2001-03) as per manufacturer’s instructions. Sequence of siRNAs used are listed in **supplementary table 2**. For inducible Pin1 knock down, HEK293 cells were transfected with lentiviruses particles containing Pin1 shRNA (Sigma-Aldrich, TRCN0000010577) as per the manufacturer’s instructions. After 24 hours, stably transfected cells were selected by treatment with 2 μg/ml puromycin (Sigma, P7255). Pin1 knock down was induced by adding 100 μM IPTG (Promega, V395A) for 48 hours. Plasmid transfections were carried out by FuGENE 6 (Roche, E269A) as per the manufacturer’s instructions.

### Ionising radiation treatment

Cells were treated with ionising radiation by CellRad Irradiator (Precision X Ray).

### Generation of GST-WW/W34A conjugated beads

Glutathione S-transferases (GST)-tagged Pin1-WW domain conjugated beads were generated as described earlier (Daza-Martin et al. 2019). Briefly, BL21 *E. coli* cells were transformed with pGEX protein expression vector containing cDNA for WW or W34A domain (obtained from JRM laboratory). Transformed colonies were later cultured in 50 ml LB media containing ampicillin for 16 hours at 37°C. The next day, 5ml of this starter culture was transferred to 500 ml fresh LB media containing ampicillin and cultured at 37°C till OD reached 0.6. At this stage, 1 mM IPTG was added to induce protein expression and cells were further cultured for 16 hours at 16°C. Cells were then pelleted and lysed in 20 ml GST lysis buffer (20 mM Tris HCl pH 8, 130 mM NaCl, 1 mM EGTA, 1.5 mM MgCl2, 1% Triton X-100, 10% glycerol, 1 mM DTT) with the addition of two EDTA-free cOmplete protease inhibitor cocktail (Roche, 11836170001). Supernatant from lysed bacteria was collected by centrifugation at 16000 g for 10 minutes at 4°C. Cleared supernatants were then incubated with 500 μl glutathione sepharose 4B beads (GE Life Sciences, 17-0756-01) overnight at 4°C. Beads were washed three times with lysis buffer and resuspended in GST storage buffer (20 mM Tris HCl pH 8, 130 mM NaCl, 10% Glycerol and 1 mM DTT) at 50% volume.

### Pull-down assay

Cells were lysed in NP40 lysis buffer (50 mM Tris HCl pH 7.4, 250 mM NaCl, 5 mM EDTA, 50 mM, 1% Nonidet P40, pH 8.0) with the addition of cOmplete protease inhibitor cocktail (Roche, 11836170001) and PhosSTOP (Roche, 04906837001) at 4°C. Cell debris was removed by centrifugation at 16000 g for 10 minutes at 4°C. Cell supernatant was then incubated with PBST (137 mM NaCl, 2.7 mM KCl, 8 mM Na_2_HPO_4_, 2 mM KH_2_PO_4_ and 1% Tween 20, pH 7.4) washed GST-WW or GST-W34A beads (20μl) for 16 hours at 4°C. Beads were then washed three times with NP40 lysis buffer and boiled at 95°C for 10 minutes in 30 μl 4x SDS loading buffer. Pull down proteins were then run on SDS-PAGE and analysed by western blotting.

### His-SUMO pull down

HEK293 cells expressing 6xHis-Flag-SUMO2 were lysed in 8M Urea buffer (8 M urea, 0.1 M Na2HPO4/NaH2PO4, 0.01 M Tris–HCl, pH 8, 10 mM β-mercaptoethanol) and sonicated at 10% intensity for 10 seconds. Cell debris was removed by centrifugation at 16000 g for 10 minutes at 4°C. Cleared supernatant was incubated with 20 μl His Mag Sepharose Ni beads (Sigma, GE28-9673-88) overnight. Beads were washed three times with PBST. Precipitated proteins were eluted by boiling beads in 30 μl 4x SDS loading buffer and analysed by western blotting.

### Western blotting

For western blotting 50 μg cell lysates or pull-down samples were denatured by addition of 4x SDS loading buffer and boiling at 95°C for 10 minutes. Proteins were separated by their molecular weight by running onto 6%, 8%, 12% SDS-PAGE gels and transferred onto PVDF membrane. The following antibodies were used in this study: rabbit anti RNF168 polyclonal Ab (1:1000 dilution, Millipore, ABE367), mouse anti myc mAb (1:1000 dilution, CST, 2276), mouse anti Pin1 mAb (1:2000 dilution, R&D Systems, MAB2294), rabbit anti Histone H3 polyclonal Ab (1:2000 dilution, Abcam, ab1791), rabbit anti γ-H2AX polyclonal Ab (1:1000 dilution, Abcam, ab2893), rabbit anti H2AX polyclonal Ab (1:1000 dilution, Abcam, ab11175), mouse anti SUMO1 mAb (1:500 dilution, Millipore, MABS2071), mouse anti SUMO2/3 mAb (1:1000 dilution, Abcam, ab81371), rabbit anti RNF8 mAb (1:1000 dilution, Abcam, ab128872) mouse anti Tubulin mAb (1:1000 dilution, Santa Cruz, sc-5286), rabbit anti Vinculin mAb (1:2000 dilution, Abcam, ab129002), rabbit anti-mouse-HRP polyclonal Ab (1:5000 dilution, DAKO, P0161), Swine anti-rabbit polyclonal Ab (1:5000 dilution, DAKO, P0217).

### Preparation of chromatin fraction

U2OS cells were seeded at 50% confluency in 10cm plates and treated with various siRNAs and 1 μg/ml doxycycline for 72 hours. Cells were harvested by trypsinisation and washed two times with PBS and resuspended in 500 μl PBS. 50 μl of cells were kept aside for the preparation of whole cell lysate (WCL). The remaining cells were resuspended in ice-cold sucrose buffer (10 mM Tris-Cl pH 7.5, 20 mM KCl, 250 mM Sucrose, 2.5 mM MgCl2, 0.3% Triton X-100, cOmplete protease inhibitor cocktail, PhosSTOP, 50 μM PR619 (Sigma, SML0430) and 20 μM MG132) and vortexed three times at low speed for 5 seconds. Cells were centrifuged at 500 g for 5 minutes at 4°C and supernatant was saved as cytoplasmic fraction. Pellet was resuspended in 200 μl NETN buffer (50 mM Tris-Cl pH 8.0, 150 mM NaCl, 2 mM EDTA, 0.5 % NP-40, cOmplete protease inhibitor cocktail, PhosSTOP, 50 μM PR619 and 20 μM MG132) and kept on ice for 30 minutes with intermittent tapping and centrifuged at 1700 g for 5 minutes at 4°C. Supernatant was saved as nuclear fraction and remaining pellet (chromatin fraction) was resuspended in 200 μl NETN buffer. Both WCL and chromatin fraction were sonicated twice at 5% intensity for 10 seconds on ice and denatured by the addition of 4x SDS loading buffer and boiling at 95°C for 5 minutes. Samples were analysed by western blotting.

### Immunofluorescence microscopy and quantification of fluorescence intensity

For immunofluorescence microscopy, 1×10^4^ cells were plated on 13 mm glass coverslips. Cells were transfected with siRNAs and 1 μg/ml doxycycline for 72 hours. For 53BP1, cells cultures were supplemented with 10 mM EdU for 30 minutes. After treatment with IR, cells were pre-extracted with 0.5% Triton X-100 in PBST for 5 minutes on ice and fixed with 4% paraformaldehyde. Cells were permeabilised by 0.5% Triton X-100 for 30 minutes at room temperature (RT). After blocking with 10% FCS, cells were incubated with primary antibodies for 1 hr, RT, except for mouse anti-myc mAb and rabbit anti-RNF168 polyclonal Ab (16 hours, 4°C). After three washes with PBST, cells were incubated with secondary antibodies for 1 hour, RT. DNA was stained with Hoechst stain (1:50000 dilution). Following antibodies were used in this study: rabbit anti RNF168 polyclonal Ab (1:500 dilution, Millipore, ABE367), mouse anti-myc mAb (1:500 dilution, Sigma, M5546), Rabbit anti-53BP1 polyclonal Ab (1:3000 dilution, Abcam, ab36823), mouse anti γ-H2AX mAb (1:2000 dilution, Abcam, ab22551), rabbit anti γ-H2AX polyclonal Ab (1:2000 dilution, Abcam, ab2893), Donkey anti-mouse-Alexa-488 polyclonal Ab (1:2000 dilution, Life technologies, A21202), Donkey anti-mouse Alexa-555 polyclonal Ab (1:2000 dilution, Life technologies, A31570), Donkey anti-rabbit Alexa-488 (1:2000 dilution, Life technologies, A21206), Donkey anti-rabbit Alexa-555 (1:2000 dilution, Life technologies, A31572). Edu was stained using Click-iT chemistry with Alexa-647-azide as per the manufacturer’s instructions. Images were acquired using a Leica DM600B microscope with an HBO lamp with a 100-W mercury short arc UV-bulb light source and four filter cubes, A4, L5, N3 and Y5, to produce excitations at wavelengths of 360, 488, 555 and 647 nm, respectively. 100x oil immersion objective was used, and images were captured for each wavelength sequentially.

For fluorescence intensity analysis, images were analyzed using ImageJ software. Region of interest (ROI) was drawn around the nucleus to calculate integrated density, area and mean fluorescence of background. Corrected total cell fluorescence (CTCF) was calculated using the following formula: CTCF = Integrated Density – (Area of selected cell X Mean fluorescence of background readings). CTCF values for each experiment were plotted using GraphPad Prism9.

### Colony survival assay

Colony survival assays were performed as described previously (Densham et al. 2016). 2×10^4^ U2OS cells were plated in 24 well cell plates. Cells were transfected with siRNAs and treated with 1μg/ml doxycycline for 72 hours. Cells were treated with indicated doses of IR before transferring them into six-well plate at limited density. Cells were cultured for 10-14 days. Colonies were stained with 0.5% crystal violet (BDH Chemicals) in 50% methanol and counted. Colony survival was calculated as the percentage change in colony formation following IR treatment compared to matched untreated cells. Each experiment is an average of three technical repeats, and the mean of three or more experiments was plotted using GraphPad Prism9.

### Statistics

All statistical analysis was performed by student t-test. p-value <0.05 were considered significant.

## Grant funding

Wellcome Trust 206343/Z/17/Z (A.S-C, AJG, MJ). We thank the Microscopy and Imaging Services at Birmingham University (MISBU) in the Tech Hub facility for microscope support and maintenance. We thank Alexandra Walker for critical reading of the manuscript.

## Declaration of Conflict of Interest

The authors declare no conflict of interest.

## Author contributions

ASC generated stable cells lines, performed immunofluorescence experiments, pull-down assay, chromatin fractionation, western blotting, colony survival assay and analyzed data. AJG and MJ performed cloning and site-directed mutagenesis. JRM and ASC wrote the paper. JRM conceptualized and directed the project.

---

**Supplementary figure 1.**
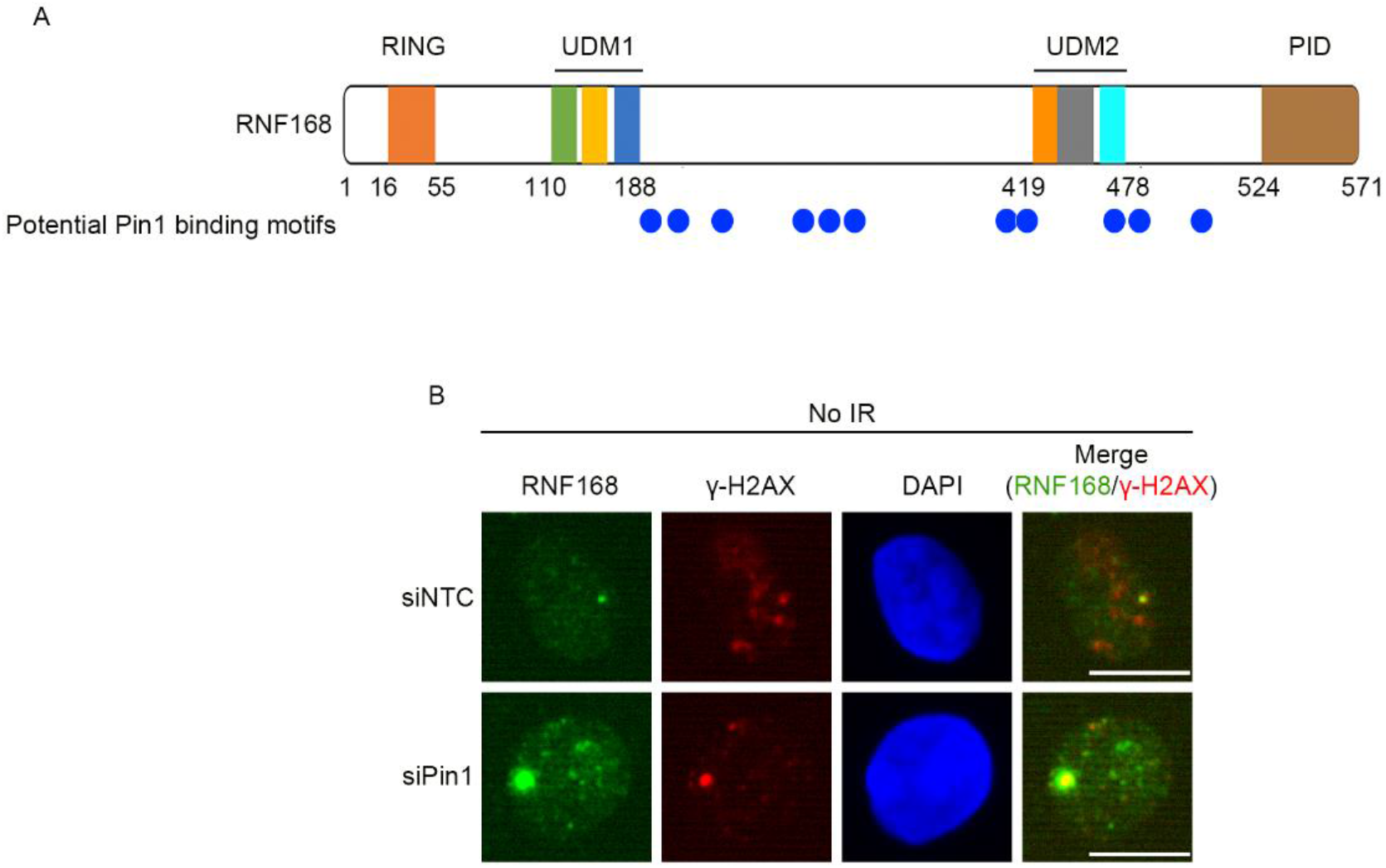
A. Diagram depicting domain architecture of RNF168. The location of potential Pin1 binding motifs in RNF168 are highlighted with blue circles. B. U2OS cells were treated with siNTC and siPin1 for 72 hrs. Later cells were fixed and stained for RNF168 and γ-H2AX. Scale bars 10 μm.

**Supplementary figure 2.**
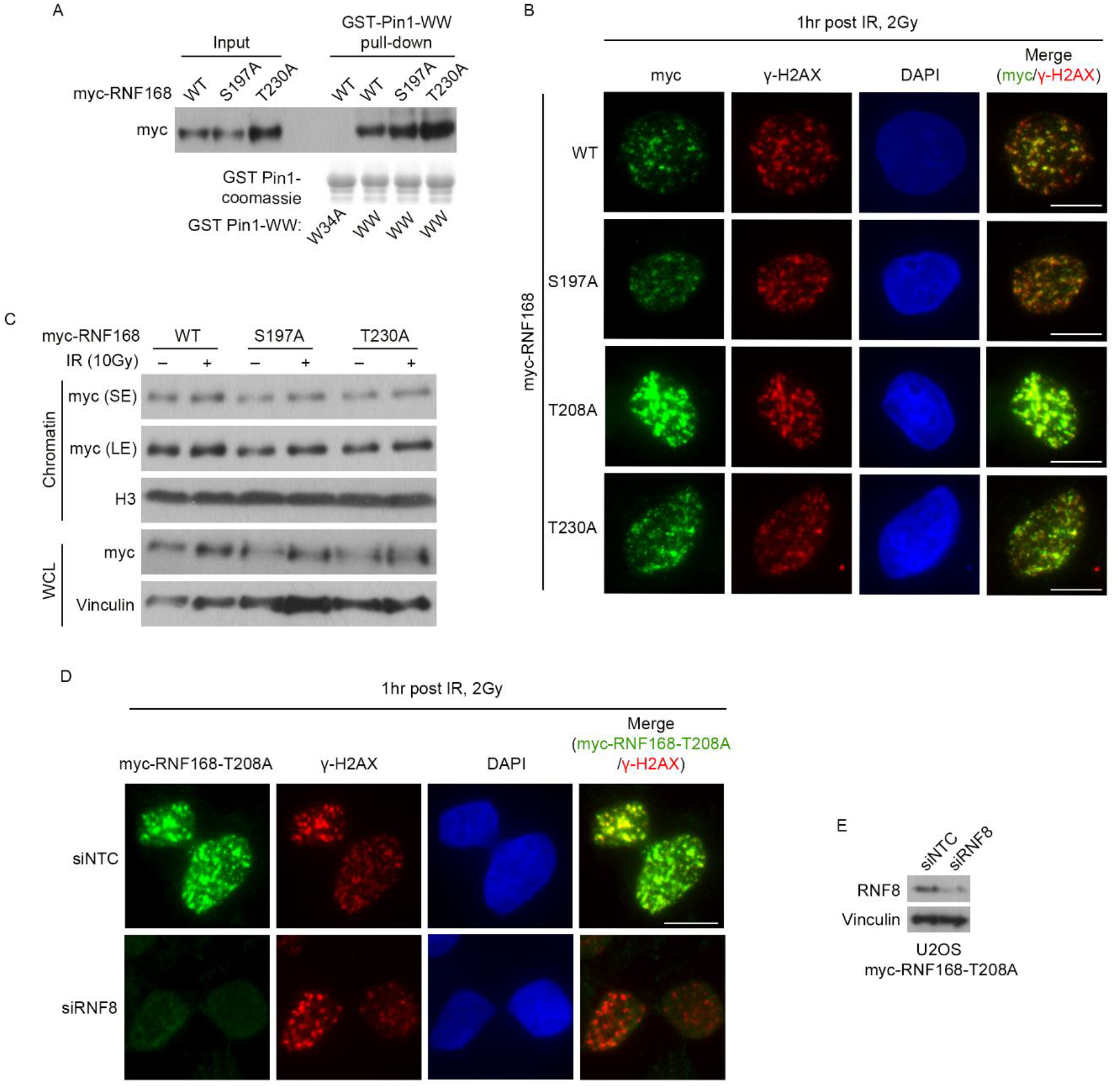
A. U2OS cells expressing myc-RNF168-WT or S197A or T230A mutant were subjected to pull down by GST-fused-WW or W34A domain of Pin1. B. U2OS cells were treated siRNF168 for 72 hrs and 1μg/ml doxycycline to induce expression of myc-RNF168-WT, S197A, T208A, T230A. Endogenous RNF168 was depleted simultaneously by treatment with siRNF168. Cells were treated with 2 Gy of IR. 1hr after IR, cells were fixed and stained for myc and γ-H2AX. Scale bars 10 μm. C. Western blot of chromatin fraction and WCL for myc, histone H3 and vinculin. Cells complemented with RNF168-WT, S197A and T230A were treated with 10 Gy of IR. 1 hr later, untreated or IR treated cells were collected to prepare chromatin and WCL fractions. D. U2OS cells complemented with myc-RNF168-T208A were treated with 1 μg/ml doxycycline and siRNF8 for 72 hrs. Cells were treated with 2 Gy of IR and fixed 1 hr later. Cells were stained for myc and γ-H2AX. Scale bars 10 μm. E. Western blot to show depletion of RNF8 for D.

**Supplementary Table 1:**
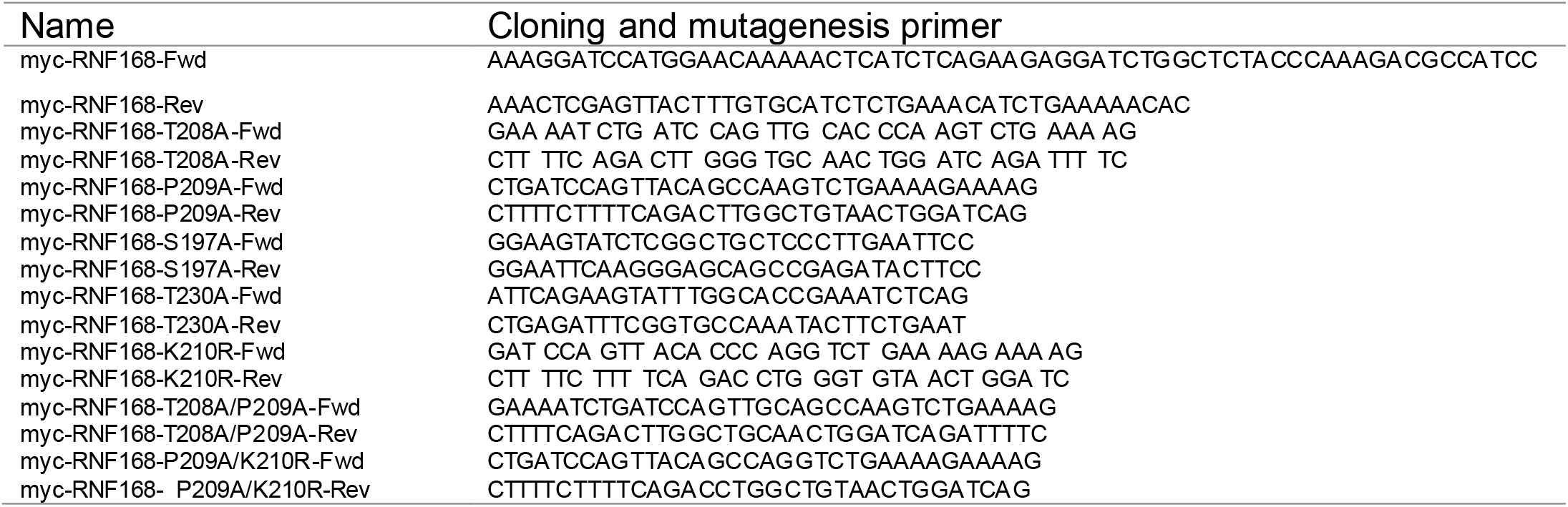
Primer sequences used for cloning.

**Supplementary Table 2:**
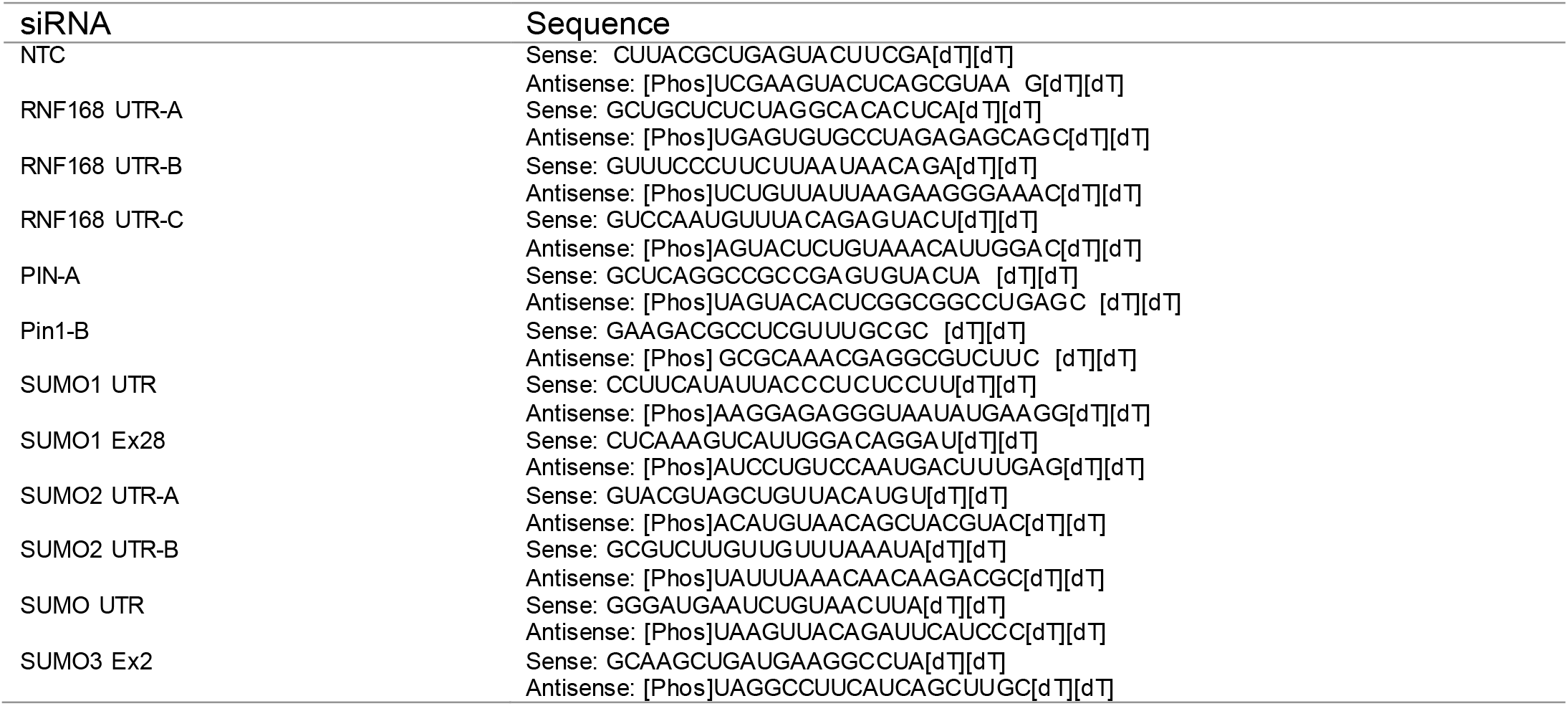
Sequences of siRNAs used.

## Notes

### Competing Interest Statement

The authors have declared no competing interest.

